# Evolution of abbreviated development in *Heliocidaris erythrogramma* dramatically re-wired the highly conserved sea urchin developmental gene regulatory network to decouple signaling center function from ultimate fate

**DOI:** 10.1101/712216

**Authors:** Allison Edgar, Maria Byrne, David R. McClay, Gregory A. Wray

**Affiliations:** Department of Biology, Duke University, Durham, NC, USA; School of Medical Science and Bosch Institute, Department of Anatomy and Histology, The University of Sydney, Sydney, NSW, Australia; School of Life and Environmental Sciences, The University of Sydney, Sydney, NSW, Australia; Center for Genomic and Computational Biology, Duke University, Durham, NC 27708, USA.

**Author notes:** **Email addresses of corresponding authors:** Allison Edgar; Gregory A. Wray). **Email addresses of co-authors:** Maria Byrne; David R. McClay.

## Abstract

Developmental gene regulatory networks (GRNs) describe the interactions among gene products that drive the differential transcriptional and cell regulatory states that pattern the embryo and specify distinct cell fates. GRNs are often deeply conserved, but whether this is the product of constraint inherent to the network structure or stabilizing selection remains unclear. We have constructed the first formal GRN for early development in *Heliocidaris erythrogramma*, a species with dramatically accelerated, direct development. This life history switch has important ecological consequences, arose rapidly, and has evolved independently many times in echinoderms, suggesting it is a product of selection. We find that *H. erythrogramma* exhibits dramatic differences in GRN topology compared with ancestral, indirect-developing sea urchins. In particular, the GRN sub-circuit that directs the early and autonomous commitment of skeletogenic cell precursors in indirect developers appears to be absent in *H. erythrogramma*, a particularly striking change in relation to both the prior conservation of this sub-circuit and the key role that these cells play ancestrally in early development as the embryonic signaling center. These results show that even highly conserved molecular mechanisms of early development can be substantially reconfigured in a relatively short evolutionary time span, suggesting that selection rather than constraint is responsible for the striking conservation of the GRN among other sea urchins.

## Introduction

Instructions encoded in the genome are executed during development to specify distinct cell types in specific spatial patterns. Developmental gene regulatory networks (GRNs) are formal models of the transcription factor cascades and cell signaling interactions that specify these diverse cell fates and distinct embryonic territories. Evolutionary changes in these processes are thought to underlie many interesting and novel phenotypes. However, two fundamental challenges to understanding how GRNs evolve are distinguishing stabilizing selection from inherent network features that promote stability and discriminating directional selection from phenotypically neutral developmental systems drift [1,2]. The sea urchin *Heliocidaris erythrogramma* is an ideal model system to explore these questions because its development has changed dramatically from the ancestral state in a relatively short evolutionary time, approximately 4 million years ago (mya). Developmental GRNs from sea urchins diverged ∼40-270 mya, and from several echinoderm outgroups diverged up to 550 mya, are particularly well-studied (reviewed in [3-6]). Thus evolution of echinoderm GRNs may be compared across orders of magnitude of divergence time.

The euechinoid genus *Heliocidaris* encompasses a dramatic shift in developmental life history [7,8]. *H. tuberculata* exhibits the ancestral condition for sea urchins: small eggs with indirect development via a feeding larva (planktotrophy). The developmental GRN underlying this ancestral life history (Figure 1A) has been characterized in considerable detail and is highly conserved across euechinoid sea urchins [6,9]. *H. erythrogramma*’s ancestors diverged from the ancestral condition, acquiring much larger eggs and greatly accelerated development via a nonfeeding larva (lecithotrophy) with highly derived morphology [8,10,11] (Figure 1B). Despite these substantive differences, the post-metamorphic phase of its life cycle is nearly indistinguishable from that of its congener *H. tuberculata*. The *Heliocidaris* lineages with ancestral and accelerated development diverged only ∼4 million years ago (mya) [7]. While this life history switch has arisen multiple times in echinoderms [8,12,13], the *Heliocidaris* genus remains the best studied example [14-33]. Loss of the feeding larval stage entails tradeoffs among maternal investment, offspring survival, and dispersal [12,34-36], although the ecological consequences of the transition to lecithotrophy are complex and incompletely understood [37]. Lecithotrophic development in *H. erythrogramma* is accompanied by dramatic changes to embryogenesis including changed timing of key developmental events, altered cleavage pattern, axial patterning, and early cell fate specification. The rapidity with which this developmental mode has arisen in *H. erythrogramma* and its implications for ecology, suggest that accelerated development is a product of strong selection for accelerated development rather than of evolutionary drift or selection on an adult trait.

**Figure 1.**
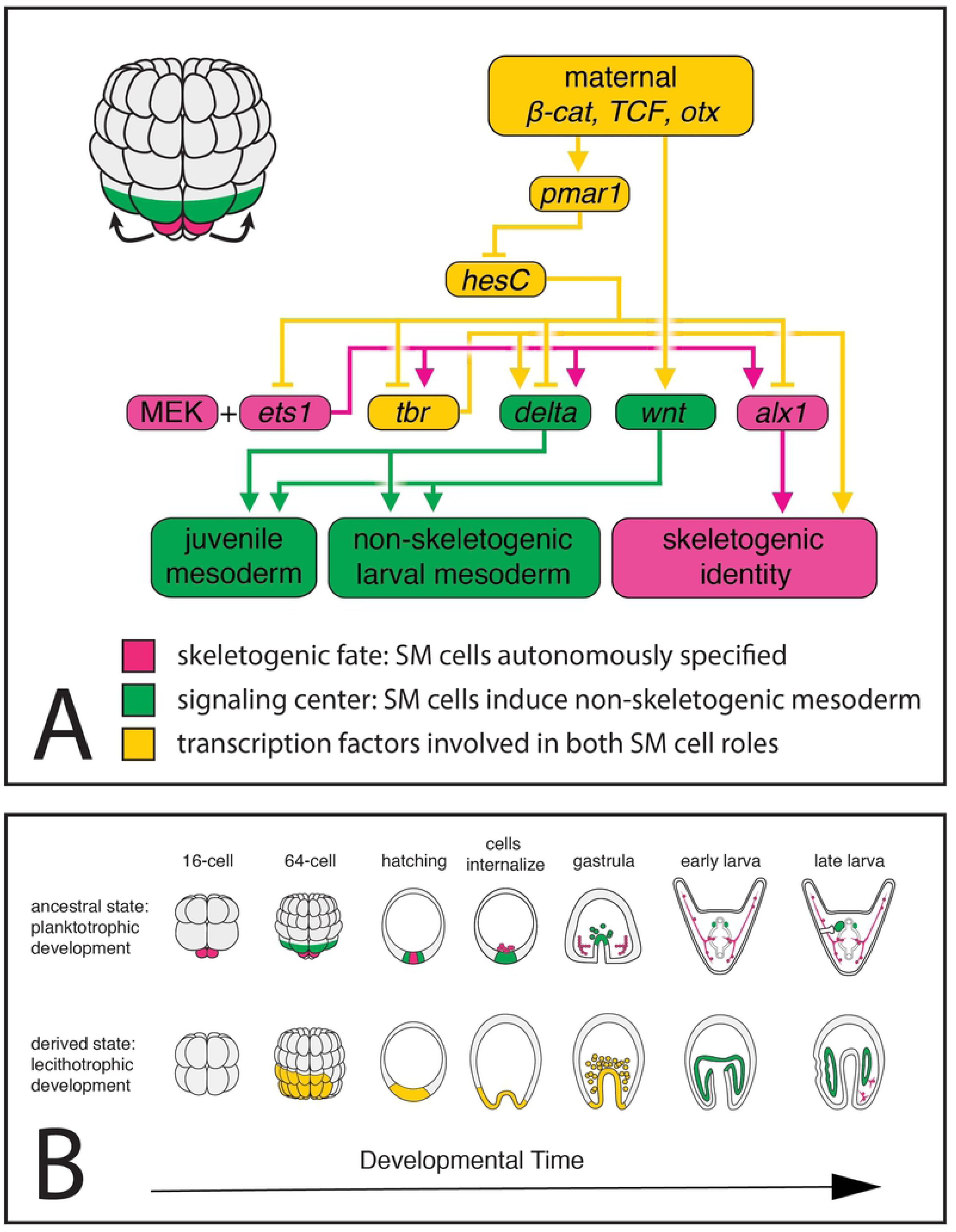
**A** Euechinoid mesodermal cell lineages are specified cell-autonomously and non-autonomously. The skeletogenic mesenchyme (SM) cells arise by an unequal cleavage (smaller pink cells) and are autonomously specified as the signaling center and prospective skeletogenic cells. A signaling relay (arrows) initiated by SM cells specifies adjacent cells as non-skeletogenic mesoderm (NSM) (green). The gene regulatory network activated in these cells was constructed by individual gene perturbation experiments to identify transcription factors and ligands that must be expressed in the SM cells for their cell-autonomous specification as prospective skeletogenic cells (pink), their identity and/or function as a signaling center (green), or both (yellow). **B** Top: Development in euechinoid planktotrophs drawn from literature. In the ancestral state, the SM cells ultimately become larval skeleton (pink), but also function as the signaling center that induces specification of other cell lineages, such as coelomic pouch mesoderm, pigment cell mesoderm and other non-skeletogenic mesenchyme (green). Coelomic pouch mesoderm is the source of juvenile skeletogenic cells (not pictured). Bottom: Model of development in the lecithotroph *H. erythrogramma* drawn from literature. The timing, lineage, and fate of the embryonic signaling center is not yet known in detail. No mesodermal sub-types are segregated before the 64-cell stage and descendants of the yellow shaded area will contribute to skeletogenic mesoderm, coelomic pouch, pigment cell mesoderm and other NSM, and endoderm. Whether and when larval (pink) and juvenile (not pictured) skeletogenic cells are segregated is unknown.

To understand how the ancestral GRN may have changed to accommodate the shift to lecithotrophy, we chose to focus on a specialized cell lineage shared by all euechinoid sea urchins [38] that is well studied for its unique developmental role, cell behaviors, and specification process: the skeletogenic mesenchyme (SM). In the planktotroph four cells specified early in development, the large micromeres which become the primary mesenchyme cells, function as a signaling center that can induce a secondary axis in the early embryo [39,40] and also are committed to become the cells that synthesize the larval endoskeleton (reviewed in [41,42].) The GRN sub-circuit responsible for specification of sea urchins’ larval SM cells, the SM-GRN, emerged >250 mya and is nearly invariant among species that diverged ∼40 mya [9]. In indirect developers distinct cell populations synthesize larval and juvenile skeletons weeks apart in development [43]. The larval skeleton is hypothesized to be a co-option of adult echinoderm endoskeleton [44,45].

*H. erythrogramma* has a greatly reduced larval skeleton and accelerated juvenile skeleton [26] but whether the GRNs underlying the larval and juvenile skeletons are conserved with the ancestral euechinoid was unknown. Prior published work on this species assumed that some or all of the mesenchyme cells that ingress into the blastocoel during gastrulation (Figure 1B) are skeletogenic, similar to the SM cells in the planktotroph that ingress shortly before gastrulation, albeit delayed (*e.g.* [16,25,46]). However, we were surprised to find that these cells do not express classic markers of the SM lineage. Our evidence is consistent instead with the hypothesis that the larval SM cell type does not exist in *H. erythrogramma*. We also examined and excluded the hypothesis that *H. erythrogramma*’s GRN is similar to the modified GRN activated in a planktotroph embryo experimentally depleted of SM cells, the replacement SM-GRN [47,48]. Instead, we found evidence that while many gene linkages remain intact, the *H. erythrogramma* GRN has several novel features, as well as features that resemble non-euechinoid urchins’ GRN [49-52]. Our results indicate a surprising degree of lability in the early developmental GRN associated with the evolution of lecithotrophy in *H. erythrogramma*.

## Results and Discussion

In classically studied, planktotrophic sea urchins, a single cell lineage that is fated to become larval skeletogenic mesenchyme also functions as an early signaling center to induce specification of other lineages. Since *H. erythrogramma* does not have obvious markers of this lineage such as asymmetric cleavage, we sought evidence of 1) the GRN circuit that specifies skeletogenic fate and 2) known phenotypes and gene expression outputs of signaling interactions coordinated by these cells. We probed the function of an early essential SM marker, Alx1 [53], as well as gene expression patterns of other SM and mesoderm markers. In the euechinoid GRN, few transcription factors are unique to the planktotrophic larval SM lineage as most are shared by other larval mesoderm or adult skeletogenic cells. We focused on markers of specifically larval SM cell identity, especially on the stages from blastula through early rudiment formation. To ask whether key signaling interactions of the SM cells are conserved, we investigated three key signaling pathways. In the euechinoid GRN, Wnt signaling initiates endomesoderm specification and a Delta signal segregates mesoderm from endoderm [54,55]; MEK-ERK signaling is then required for mesoderm specification to progress [56,57]. In order to understand how these signaling pathways operate in *H. erythrogramma*, we focused on two time points: hatched blastula, when multiple types of mesoderm have been specified in indirect developers but before morphogenesis, and the late larval stage when many differentiated cell types are present.

### *Alx1* is expressed but not localized to mesenchymal cells in *H. erythrogramma*

We first asked where the essential skeletogenic regulator gene Alx1 is expressed in *H. erythrogramma*, and found that its expression pattern differs from the consensus planktotroph. In the ancestral urchin GRN, Alx1 is necessary to specify skeletogenic cell fate and is expressed continuously in all and only larval SM cells from early specification through differentiation [53] and is expressed in larval and juvenile skeletogenic cells across echinoderms [45,51,52,58-60], suggesting that its skeletogenic function is deeply conserved.

We found *alx1* expressed throughout the vegetal pole at hatched blastula stage (Figure 2). Although both indirect developers and *H. erythrogramma* show expression at the extreme vegetal pole, *H. erythrogramma* shows broader expression of *alx1* at this stage and its localization pattern diverges markedly from the ancestral GRN from this point onwards. Instead of ingressing prior to gastrulation, *alx1*-positive cells remain in the archenteron during *H. erythrogramma* gastrulation. Previous work in planktotrophs has shown that *alx1* is involved in the epithelial-to-mesenchymal transition (EMT) that allows SM cells to ingress [53,61,62], while *ets1/2* is required for EMT in all mesenchymal cell types, SM and NSM [63].

**Figure 2.**
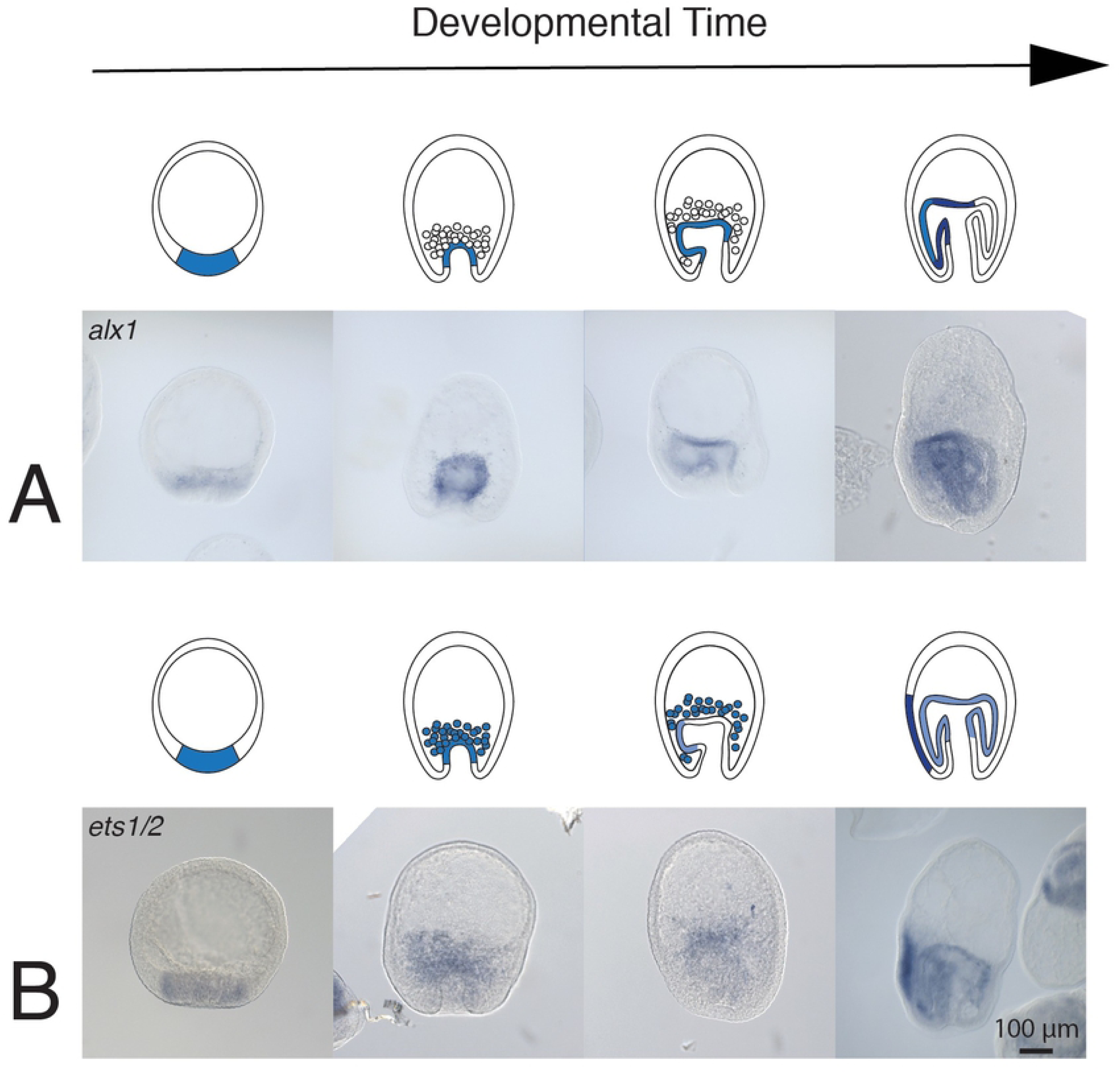
Expression of *alx1* and *ets1/2* in *H. erythrogramma*. **A** *Alx1* is expressed in the vegetal plate and archenteron throughout gastrulation, but not in ingressed mesenchyme. **B** *Ets1/2* is expressed in the vegetal plate at blastula stage, and broadly at the tip of the archenteron and ingressed mesenchyme during gastrulation, and throughout the both coelomic pouches and in the specialized ectoderm that will contribute to the juvenile.

In *H. erythrogramma*, *ets1/2*-positive mesenchyme cells ingress from the archenteron but no *alx1*-positive cells are present in the blastocoel during and after this ingression. If the *alx1*-expressing cells in *H. erythrogramma* undergo EMT it is greatly delayed relative to the onset of *alx1* expression, and if the *ets1/2*-expressing cells that ingress during gastrulation later express *alx1* that expression is greatly delayed relative to EMT. During late gastrula and early rudiment stages *alx1* is expressed throughout the region homologous to the left coelomic pouch whereas *ets1/2*-expressing cells are apparent throughout the prospective juvenile in both coelomic pouches and in the vestibular ectoderm. In later stages *alx1* is expressed in juvenile skeletogenic centers (Supplemental Figure 1), as in other echinoderms [45,59].

In all the diverse echinoderm classes known to produce larval skeleton, some or all of the *alx1*-expressing cells ingress into the blastocoel before or during gastrulation, where they continue to express *alx1* [51-53,56,60,64-67]. However, an Alx gene is also expressed in the embryos of echinoderms that do not produce larval skeletons: an alternative spliceoform of *alx1* (or a closely related paralog, Alx4/Calx) is present in the vegetal plate and mesodermal bulb of sea stars [60,66,68]. Its function there is unknown, raising the possibility that Alx1 has an alternative or additional function at the vegetal pole or in mesoderm specification. Therefore, we decided to examine the function of Alx1 in *H. erythrogramma*.

### Alx1 is necessary for skeletogenesis in *H. erythrogramma*

We used a translation-blocking morpholine-substituted antisense oligonucleotide (MASO) specific to *alx1* to examine its function *in vivo*. While Alx1 morphants are delayed overall in indirect developers, eventually all other larval mesoderm sub-types are recovered through regulative processes so the knockdown phenotype is specific to SM cells [53]. We found that Alx1 is indeed required for biomineralization of the skeleton in *H. erythrogramma*. Blocking Alx1 translation eliminates both larval and juvenile spicules (Figure 3) but does not eliminate any other cell lineage, just as in indirect developers. However, we did notice a secondary, unexpected phenotype in Alx1 morphants: the primary body axis is shortened (mean decrease 15.5% body length, two-sample t (17) = 2.136, p = 0.023; raw data in Supplemental File 1).

**Figure 3.**
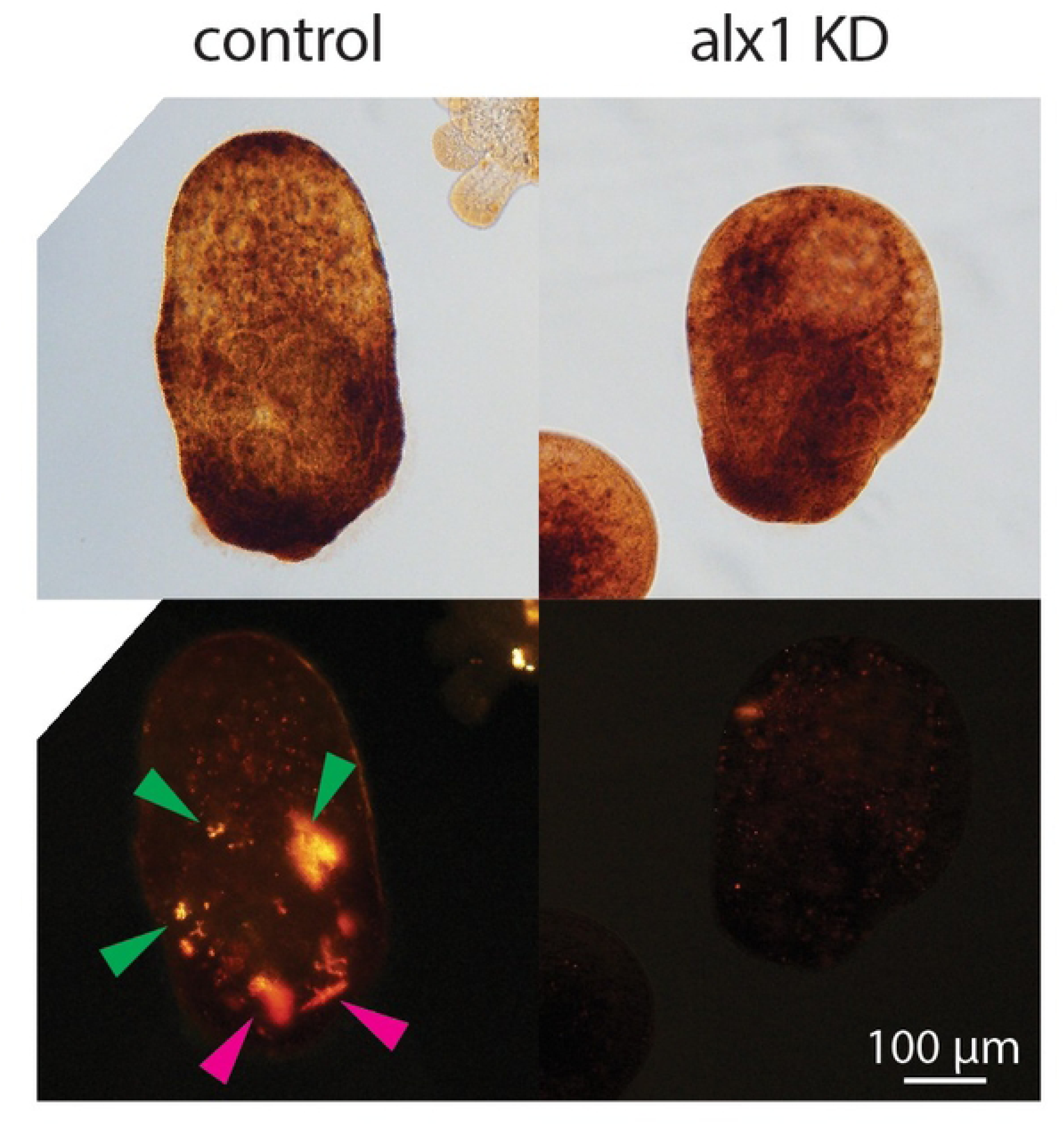
Alx1 functions in skeletogenesis in *H. erythrogramma* despite its absence in ingressed mesenchyme. DIC and polarized light views of standard control and *alx1* translation-blocking morpholino injected *H. erythrogramma*. Skeletogenesis is impaired in *alx1* morphants.

Thus, Alx1 appears to retain a skeletogenic function in *H. erythrogramma*. Since the key skeletogenic marker *alx1* and the key mesenchyme marker *ets1/2* are not co-expressed as in planktotrophs, we next examined other markers of SM cells to ask whether they were co-expressed with *alx1* to test the hypothesis that SM cell identity was maintained but EMT bypassed or delayed.

### Key genes of the ancestral larval SM-GRN are not co-expressed in *H. erythrogramma*

Like Alx1, most other SM-GRN genes are also expressed in both larval and adult skeletogenic cells of indirect developers [45]; very few genes are uniquely expressed in larval SM cells and known to be absent in juvenile and adult skeletogenic cells or other larval mesoderm. Thus, co-expression of a suite of transcription factors is the best current diagnostic marker of the euechinoid larval skeletogenic lineage.

We found that components of the larval SM-GRN do not mark a single persistent cell population in *H. erythrogramma* as in indirect developers and no group of cells co-expresses the genes of the ancestral larval SM-GRN after blastula stage. Neither the *ets1/2*-positive mesenchyme nor the *alx1*-positive coelomic pouch mesoderm co-express key diagnostic SM-GRN genes, so it is not simply that one gene was lost from the conserved sub-circuit (or failure of a single probe). Low sequence divergence between *H. erythrogramma* and a closely related congeneric species, *H. tuberculata*, permits probe hybridization across species under identical hybridization conditions. We used this to test the hypothesis that changes in the expression pattern between *H. erythrogramma* and the ancestral GRN arose concurrently with accelerated development rather than as a difference in the Heliocidaris lineage from other planktotrophic sea urchins where the expression of these genes is well characterized. The T-box gene Tbr was restricted to the SM lineage in euechinoid urchins [51,69] rather than its ancestral role in pan-mesodermal and broad endomesoderm specification [38,60,65,70,71] but it remains indispensable to activate the normal endomesoderm GRN [72,73] and the replacement SM-GRN [47,74]. *Tbr*’s placement in the GRN immediately downstream of the HesC/Pmar1 logic gate and integration into a circuit with Alx1 has been proposed as the key event in the evolution of the larval SM cell type [38,45,75]. Thus, Tbr is a key node that integrates the cell identity and signaling center functions of the sea urchin micromere lineage.

Our data suggest that this Ets1/2-Alx1-Tbr sub-circuit is absent or transient in *H. erythrogramma*. All three genes show distinct spatiotemporal expression patterns rather than co-expression in *H. erythrogramma*. Whereas the expression patterns of *ets1/2* and *tbr* in *H. tuberculata* resemble closely patterns seen in other planktotrophs, in *H. erythrogramma tbr* expression is lost at the onset of gastrulation and is not seen in mesenchyme (Figure 4). Despite the different physical localization of *tbr* transcripts in planktotrophs and lecithotrophs (Figure 4B), whole-transcriptome temporal expression of *tbr* and *alx1* do not differ (Supplemental Figure 2). Like Tbr, FoxB is expressed in both the normal [72] and replacement SM-GRNs [47] but not in the juvenile skeletogenic cells [45], and was likely co-opted into this GRN in the lineage leading to urchins as it is absent from brittle star larval SM cells [65]. In *H. tuberculata foxB* is expressed in the skeletogenic mesenchyme cells as in other planktotrophs but *foxB* is not expressed in either the *ets1/2*-positive mesenchyme or in the *alx1*-positive territory of the archenteron in *H. erythrogramma* (Figure 4C). *FoxB* is expressed in *H. erythrogramma*’s later larval stages (Supplemental Figure 2) but not in the skeletogenic centers (not shown).

**Figure 4.**
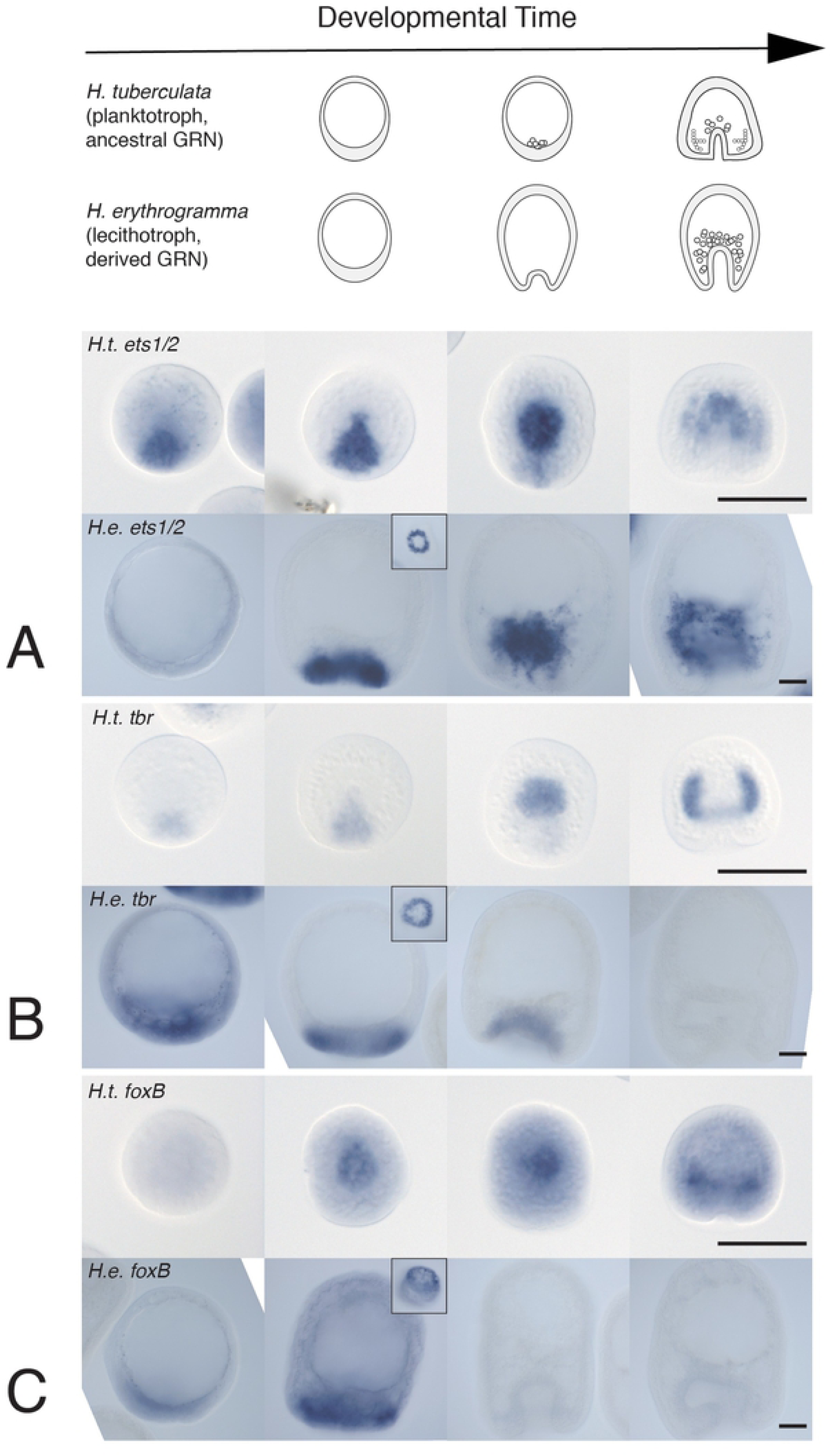
Expression of key larval SM marker genes in *H. tuberculata* (top rows) and *H. erythrogramma* (bottom rows) at equivalent stages. Note that the two species are different sizes; both scale bars represent 50 μm. Insets show vegetal views. All three genes (*ets1/2, tbr, foxB*) are known to be expressed in the replacement SM-GRN as well as the normal SM-GRN. **A** *Ets1/2* is expressed in many different mesoderm sub-types in *H. tuberculata* as in other planktotrophs. **B** *Tbr* is expressed exclusively in larval SM cells in the ancestral SM-GRN and this is conserved in *H. tuberculata*. *Tbr* shows a different expression pattern in *H. erythrogramma*; *tbr* is expressed similarly to the ancestral pattern at early stages but is not found in mesenchymal cells at any stage. At blastula, *tbr* is expressed in an asymmetric ring at vegetal pole. At early gastrula; *tbr* is expressed in the invaginating archenteron but not in the early ingressing mesenchyme; localized *tbr* expression is not seen after this time point. **C** *FoxB* is expressed in *H. tuberculata* similarly to other planktotrophs, in SM cells (as well as the archenteron and ventral ectoderm in later stages, not shown). *FoxB* is co-expressed with *ets1/2* and *tbr* in *H. erythrogramma*’s vegetal pole at blastula stages but is not found in mesenchymal cells at any stage. *FoxB* expression is lost in *H. erythrogramma* after onset of gastrulation.

We also examined other members of the SM-GRN. We found no evidence of localized expression for the SM lineage-specific repressor *pmar1* [76,77] in *H. erythrogramma* (Figure 5A). Pmar1 paralogs appear to have duplicated repeatedly and diversified independently in various euechinoid species [78] so it is possible that we have not identified the functionally relevant paralog. However, our *pmar1* probe shows specific expression in *H. tuberculata* SM cells. We also found that the endomesodermal Forkhead transcription factor *foxN2/3* is expressed similarly in *H. erythrogramma* as in the ancestral euechinoid (Figure 5B). In the ancestral euechinoid GRN, *foxN2/3* is found in pre-ingression SM; its later expression shifts to other endomesodermal territories [79,80], similar to the pattern in *H. erythrogramma*. Thus, *foxN2/3*’s expression pattern is consistent with a role in endomesoderm specification.

**Figure 5.**
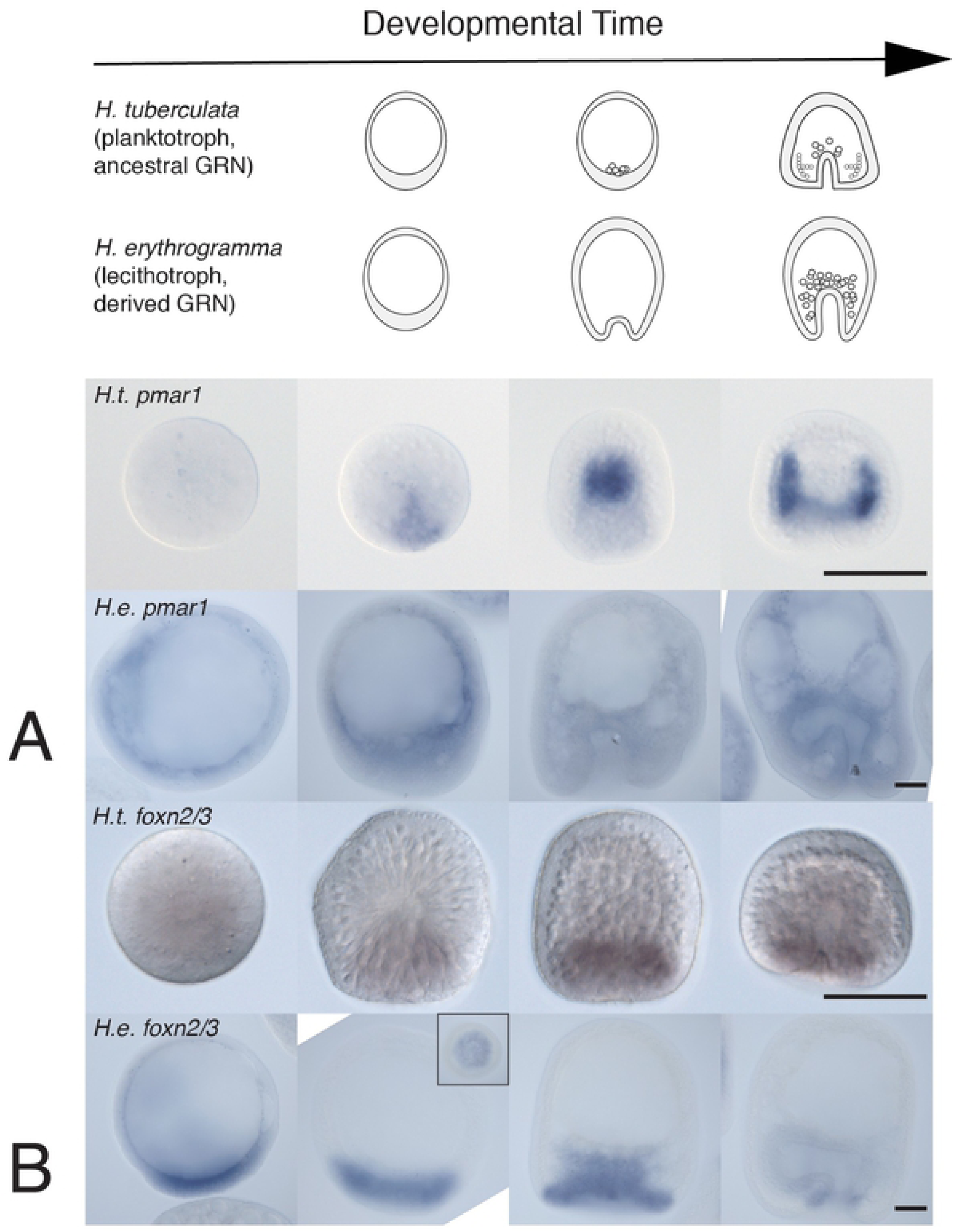
Expression of other larval SM marker genes in *H. tuberculata* (top rows) and *H. erythrogramma* (bottom rows) at equivalent stages. Note that the two species are different sizes; both scale bars represent 50 μm. **A** *Pmar1* is expressed only in the normal SM-GRN in planktotrophs, not the replacement SM-GRN. We were not able to detect *pmar1* expression in *H. erythrogramma* although the probe shows expression in *H. tuberculata* SM cells. Nonspecific chromogenic staining is visible inside the blastocoel. **B** In both *H. tuberculata* and *H. erythrogramma*, *foxN2/3* is expressed similarly to the consensus euechinoid. *FoxN2/3* is expressed similarly at the vegetal plate and at the tip of the archenteron, but not in fully ingressed mesenchyme; later it is expressed in the hindgut. Inset shows vegetal view.

There is a brief window at hatched blastula stage in which the characteristic SM-GRN genes *alx1*, *ets1/2*, *tbr*, *foxB*, and *foxN2/3* are co-expressed in the vegetal plate but they never again show co-expression in any *H. erythrogramma* cell type. Expression of SM differentiation genes downstream of these early genes [81] is absent, reduced, or delayed relative to indirect developers (Supplemental Figure 2). Taken together, these data suggest that the larval SM cell lineage known from indirect developers has been lost from *H. erythrogramma*. We next considered whether signaling functions coordinated by SM cells in indirect developers were altered in *H. erythrogramma* and found some striking differences.

### Early canonical Wnt signaling activates mesodermal genes differently in *H. erythrogramma* than in indirect developers

Canonical Wnt (cWnt) signaling is a deeply conserved activator of endomesodermal development across bilaterians, including sea urchins [82-84]. The ancestral GRN predicts that cWnt signaling should expand endoderm at the expense of ectoderm without dramatically affecting mesoderm. However, it is thought that early endomesoderm fate specification does not require a secreted Wnt signal but instead nuclearization of maternally loaded β-catenin [85]. Activation of cWnt with the GSK3-β inhibitor LiCl does not expand expression domains of the mesoderm markers *delta* and *tbr* in indirect developers [55]. Reciprocally, in the indirect developer *S. purpuratus*, treatment with the PORCN inhibitor C59, which prevents secretion of Wnt ligands, does not affect expression levels of the key mesodermal genes Alx1, Ets1/2, Tbr, or Gcm (<0.2 fold-change [86]).

Previous work in *H. erythrogramma* showed that activation of cWnt causes exogastrulation [32]. Axin and GSK3-β work together to destabilize β-catenin, an effector of cWnt signaling. We found that a translation-blocking MASO targeting *axin2* phenocopies GSK3-β inhibitors, causing exogastrulation (Figure 6A, B). Reciprocally, treatment with C59 reduces the length of the archenteron. However, cWnt and GSK3-β inhibitors affect *H. erythrogramma* gene expression differently than the ancestral GRN (Figure 6C-H).

**Figure 6.**
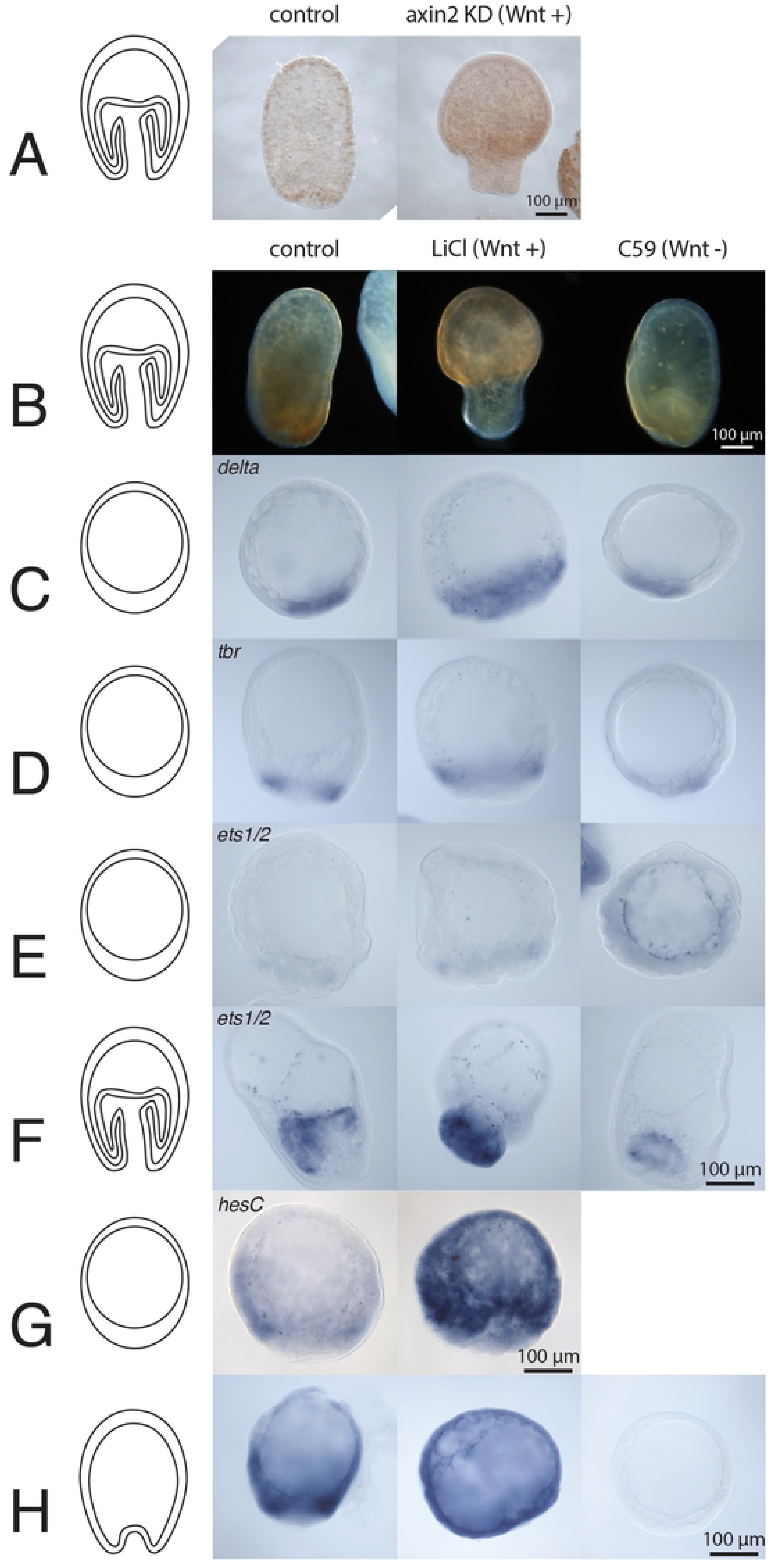
Outputs of the canonical Wnt signaling pathway in *H. erythrogramma* differ from predictions of the ancestral euechinoid GRN. **A** A translation-blocking morpholino targeting *axin2* induces exogastrulation. **B** The GSK3-β inhibitor LiCl induces exogastrulation and and the PORCN inhibitor C59 reduces the archenteron. **C-H** Mesoderm and SM marker gene expression patterns in Wnt pathway perturbed *H. erythrogramma* blastulae. Insets show vegetal views.

In *H. erythrogramma*, GSK3-β inhibitor treatment expands expression of *delta*, a marker for SM and NSM in indirect developers, throughout the vegetal pole (Figure 6C). This is unlike LiCl-treated planktotrophs, which have an essentially normal *delta* expression pattern [55]. While some treated *H. erythrogramma* embryos show a slight shift of the *ets1/2* and *tbr* expression domains towards the animal pole, their expression does not expand towards the vegetal pole as *delta* does with LiCl treatment (Figure 6D, E). This result is similar to what is seen in planktotrophs [55], but in the context of *delta* expansion suggests that *delta*, *ets1/2*, and *tbr* are not tightly co-regulated as they are as in the ancestral GRN.

Later, during gastrula stages, Wnt signaling dramatically affects mesoderm in *H. erythrogramma* as *ets1/2* expression is expanded in LiCl-treated embryos and reduced in C59-treated embryos (Figure 6F). *Ets1/2* expression in the exogastrulated cells is consistent with the observation that much of the archenteron is coelomic pouch mesoderm rather than endoderm in *H. erythrogramma* [87]. The C59 results show that *ets1/2* expression likely requires a secreted Wnt signal in *H. erythrogramma*, suggesting that *ets1/2* transcription is initiated by a GRN that resembles the ancestral endomesoderm GRN, not the SM-GRN.

In the ancestral euechinoid GRN, *hesC* is repressed downstream of cWnt (by endogenous Tcf/β-catenin via Pmar1) and thus the increased expression of *hesC* in LiCl-treated *H. erythrogramma* was not predicted by the ancestral euechinoid GRN. While very early *hesC* expression is uniformly distributed throughout all cells except the SM in planktotrophs [75], at blastula stages and beyond its expression pattern is much more complex [88]. However, C59 only slightly decreases *hesC* expression in the planktotroph at blastula stages and beyond (<0.1-0.3 fold-change [86]), confirming that cWnt is not a major regulator of *hesC* in the ancestral GRN. However, in *H. erythrogramma*, we find that cWnt is a major driver of *hesC* expression at these stages.

### Skeletogenic and non-skeletogenic mesenchyme are specified independently of Delta-Notch signaling in *H. erythrogramma*

To investigate another key ancestral pathway, we focused on Delta-Notch signaling. In indirect developers a Delta signal from SM cells induces specification of non-skeletogenic mesoderm (NSM) [89-91]. Perturbing Delta-Notch signaling during different critical periods eliminates distinct mesodermal cell populations such as pigment cells and coelomic pouch (which gives rise to adult structures) [92,93]. We found that two populations of mesoderm respond similarly to Delta inhibition in *H. erythrogramma* as in the ancestral GRN but one population is regulated differently.

We inhibited Delta signaling in *H. erythrogramma* by preventing translation of *delta* mRNA with an injected MASO or preventing cleavage of the Notch intracellular domain by treatment with gamma-secretase inhibitors. High and low doses of MASO or inhibitor abrogated or reduced coelomic pouch formation at all time points tested (Figure 7, Supplemental Figure 3A), just as in the ancestral GRN. At high doses of MASO (Figure 7A) or inhibitor (Supplemental Figure 3D), axial patterning and gastrulation are disrupted. At low doses of either the inhibitor or MASO, although gastrulation is abnormal some endoderm is internalized and differentiates (Figure 7B,C).

**Figure 7.**
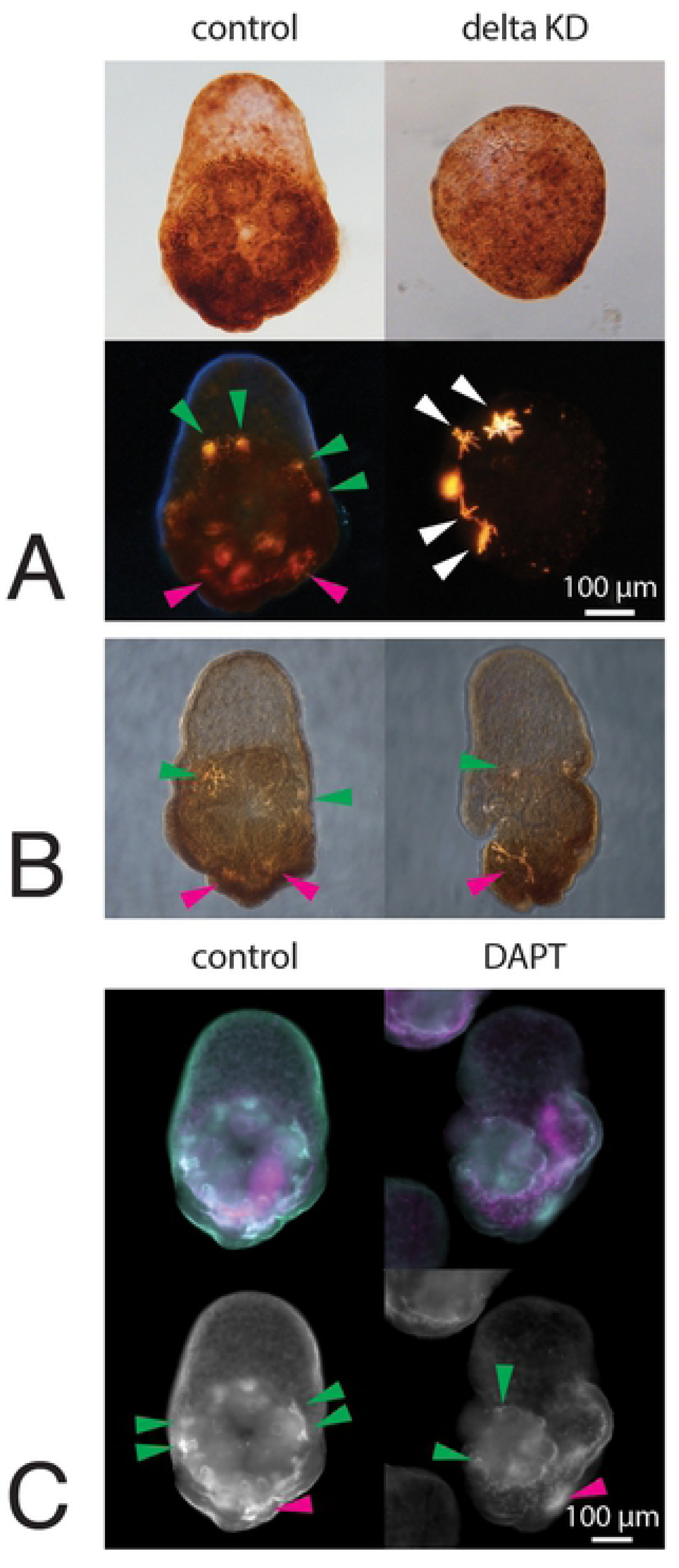
The cell types affected by disrupting Delta-Notch signaling in *H. erythrogramma* overlap with but differ from known planktotroph phenotypes suggesting that the role of Delta signaling has changed from the ancestral GRN. **A** Control morpholino-injected embryos show pentameral patterning and red pigmentation under white light (top) and both larval and juvenile skeleton under polarized light (bottom). Larval (pink arrowhead shows example) and juvenile (green arrowheads show examples) skeletal elements can be distinguished morphologically. High dose *delta*-targeting morpholino radializes the embryo but does not eliminate pigmentation or skeleton (white arrowheads); lack of normal morphological markers prevents assignment of skeleton as larval or juvenile. Late larval stage, oral view. **B** Low dose *delta*-targeting morpholino permits gastrulation but reduces juvenile rudiment size. Both larval (pink arrowheads) and juvenile (green arrowheads) skeleton can be distinguished morphologically. **C** Low-dose DAPT treatment produces a similar phenotype to low-dose morpholino injection; differentiated skeletogenic cells (skeletal marker msp130 antibody 1D5, green) and endoderm marker (EndoI, magenta) are present. Skeletal marker single channel shown below; larval (pink arrowheads) and juvenile (green arrowheads) skeleton can be distinguished morphologically. Additional skeletogenic cells are visible scattered especially in the ectoderm in and near the vestibule and juvenile skeleton morphogenesis is abnormal.

Also as in indirect developers, *H. erythrogramma* do not require a Delta signal to specify skeletogenic cells. Delta is never necessary for skeletogenic cell fate specification in the ancestral GRN. Even when SM cells are experimentally depleted, the alternative mechanism by which they are replaced (the replacement SM-GRN) does not require Delta [47]. Even in the absence of a normal rudiment, *H. erythrogramma* skeletogenic cells differentiate and respond to ectodermal patterning cues by migrating to the normal location of larval skeleton and the ectoderm region which normally would contribute to the juvenile (Figure 7C, Supplemental Figure 3C). Thus, Delta signaling is required for specification of coelomic pouch cells but not skeletogenic mesenchyme *H. erythrogramma*, just as in the ancestral GRN.

In contrast, the requirement for Delta signaling in pigment cell fate specification in the ancestral GRN appears to be lost in *H. erythrogramma*. The ancestral euechinoid specification of pigment cells requires Delta [89,91-95] and no regulative mechanism replaces this cell type if the Delta signal is absent during the early critical window, while the other mesodermal lineages can be replaced [47,48]. Even *H. erythrogramma* embryos exposed to high doses of morpholino or drug contain abundant pigment cells (Figure 7A; Supplemental Figure 3D). We did not observe a delay in the appearance of pigmentation relative to controls.

While it is not possible to conclude from the presence of both differentiated skeleton and pigment cells in Delta-perturbed *H. erythrogramma* whether these cells arose by an alternative GRN than those cell types normally do in unperturbed *H. erythrogramma*, the presence of pigment cells is a striking departure from the ancestral euechinoid GRN. These results, together with previous evidence from *H. erythrogramma*, suggest that the signaling event has been lost rather than a novel regulative mechanism gained. *H. erythrogramma*’s pigment and skeletogenic cells derive from a common lineage until at least the 64-cell stage [21,87] but potential to give rise to pigment cells is segregated by the 2-cell stage [30]. We found reduced maternal loading of *ets1/2* transcripts and dramatically increased maternal loading of the early pigment cell marker *gcm* transcripts in *H. erythrogramma* compared to planktotrophs (Supplemental Figure 2).

### Blastula-stage *H. erythrogramma* embryos have different transcription factor outputs of Delta signaling than the ancestral GRN

Next, we investigated how Delta signaling influences downstream gene expression. We found that most mesodermal genes respond differently to Delta signaling in *H. erythrogramma* than in the ancestral euechinoid. At blastula stage, DAPT-treated *H. erythrogramma* show expanded *delta* expression in the animal pole domain. This result differs dramatically from the ancestral GRN, where DAPT treatment decreases *delta* expression dramatically at the vegetal pole but does not alter the expression pattern at the animal pole [92]. A second apparent difference concerns *hesC*, which encodes a transcriptional repressor that appears to have an ancient role in segregating SM from NSM cells that predates the consensus euechinoid GRN although many of its targets are specific to euechinoids [96,97] (Figure 1).

DAPT treatment reduces *hesC* expression at both the animal and vegetal poles in *H. erythrogramma* (Figure 8). This result suggests that *hesC* expression in *H. erythrogramma* is controlled at least in part by Delta signaling. In the consensus euechinoid GRN *hesC* is usually considered to be broadly expressed upstream of *delta* [75,98], but other data suggest that delta expression precedes *hesC*’s clearance from the vegetal pole [99]. In either case, a negative regulatory relationship between Delta and HesC appears to be a euechinoid trait, as in cidaroid urchins *hesC* expression is activated at least in part by Delta [51], similar to our results in *H. erythrogramma*.

**Figure 8.**
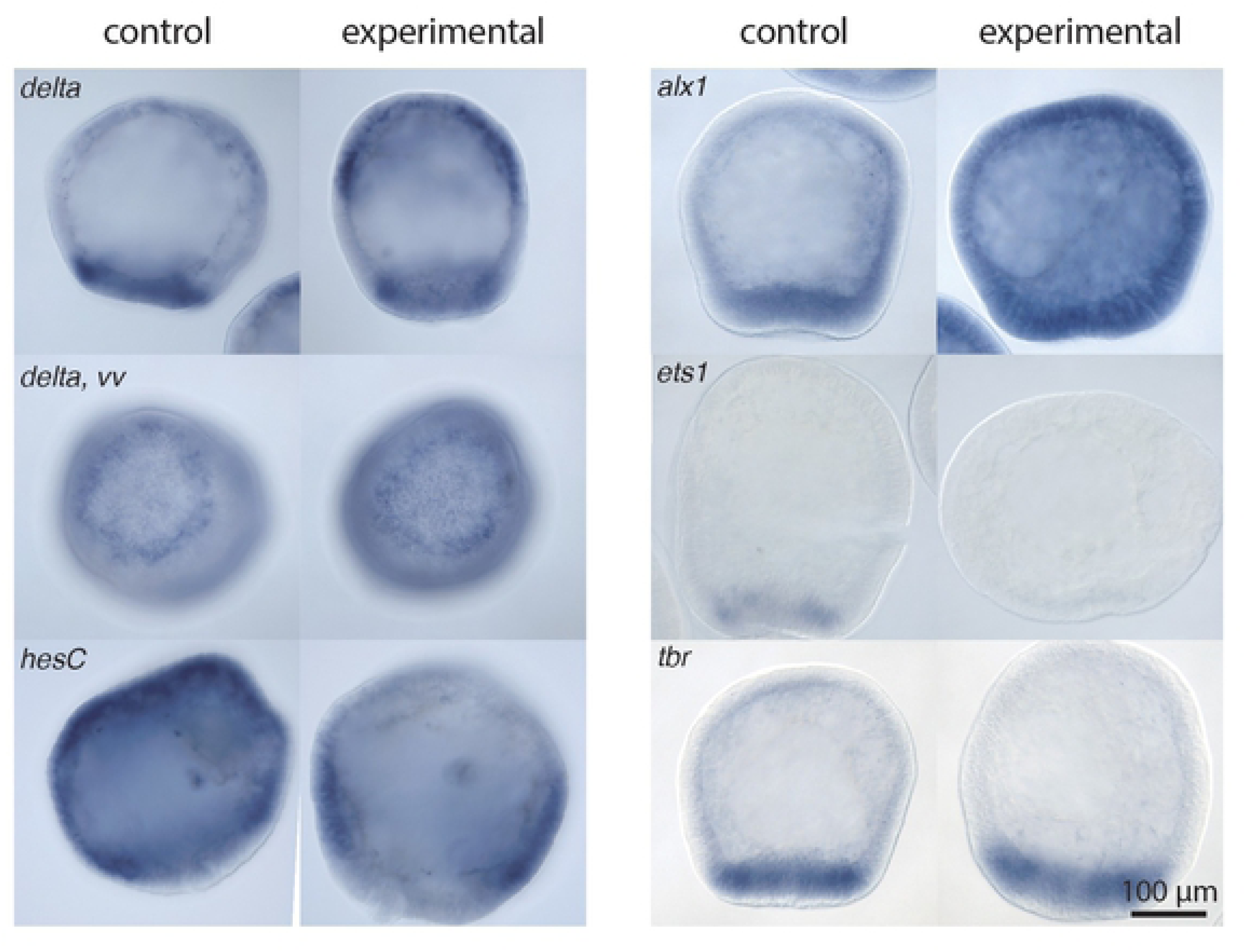
Blastula-stage gene regulatory outputs of Delta signaling in the accelerated development of *H. erythrogramma*. Interfering with Delta-Notch signaling (by treatment with the gamma-secretase inhibitor DAPT) expands *delta* expression at the animal pole without altering the vegetal pole domain expression pattern. *HesC* shows a reciprocal reduction with *delta* expansion. *Alx1* expresion is increased while *ets1/2* expression is decreased at this stage. Of these key mesoderm and SM markers, only *tbr* remains unaffected in DAPT-treated *H. erythrogramma*.

A Delta-independent positive feedback loop between the transcription factors Alx1, Ets1/2, and Tbr is characteristic of both normal and replacement SM cells in the consensus euechinoid GRN. Similarly, in *H. erythrogramma* DAPT treatment does not affect the expression of *alx1* or *tbr*; in contrast, however, a Delta signaling input appears to be required for the early phase of the pan-mesodermal marker *ets1/2* expression (Figure 8). *Ets1/2* expression later recovers (not shown). However, this dramatic difference in the initiation of early *ets1/2* expression suggests that this key mesodermal gene is not expressed early and cell-autonomously as in the ancestral GRN.

### Similar cell types require MEK-ERK cascade in ancestral and accelerated GRNs, but early transcription factor expression differs

In the ancestral sea urchin GRN, MEK-ERK signaling is required in SM cells for skeletogenic identity but not endomesodermal signaling center function [56,57]. The selective MEK inhibitor UO126 arrests SM differentiation at the time of treatment [56] and is used commonly in sea urchins to produce this phenotype [61,100]. Other larval mesoderm types also require MEK signaling; the UO126 phenotype is well characterized in the consensus GRN and thought to be mediated by preventing phosphorylation of the pan-mesodermal transcription factor Ets1/2 [56,101], which is also required to activate the replacement SM-GRN when normal SM cells are experimentally depleted (at least in part by activating *tbr* transcription) [47,102].

Like the ancestral GRN, UO126-treated *H. erythrogramma* embryos have greatly reduced skeleton and pigmentation (Figure 9). Left-right patterning within the endomesoderm but not the ectoderm is disrupted in the ancestral GRN [100] and similarly in *H. erythrogramma* the ectoderm is patterned normally along this axis although the rudiment is abnormal. However, as with Delta signaling, the similar downstream phenotype apparently conceals an alternate GRN topology as the expression of key mesodermal transcription factors at hatched blastula stage differs dramatically from the ancestral GRN. UO126 treatment eliminates the *ets1/2* expression pattern but does not affect the expression pattern of *tbr*. This is just the opposite of the ancestral GRN in which loss of *tbr* expression is diagnostic for the SM-GRN’s failure in the absence of MEK-ERK signaling [47,102] (although *tbr* expression may recover by gastrula stage in planktotrophs [61]). Later *ets1/2* expression is not affected by UO126 treatment (not shown).

**Figure 9.**
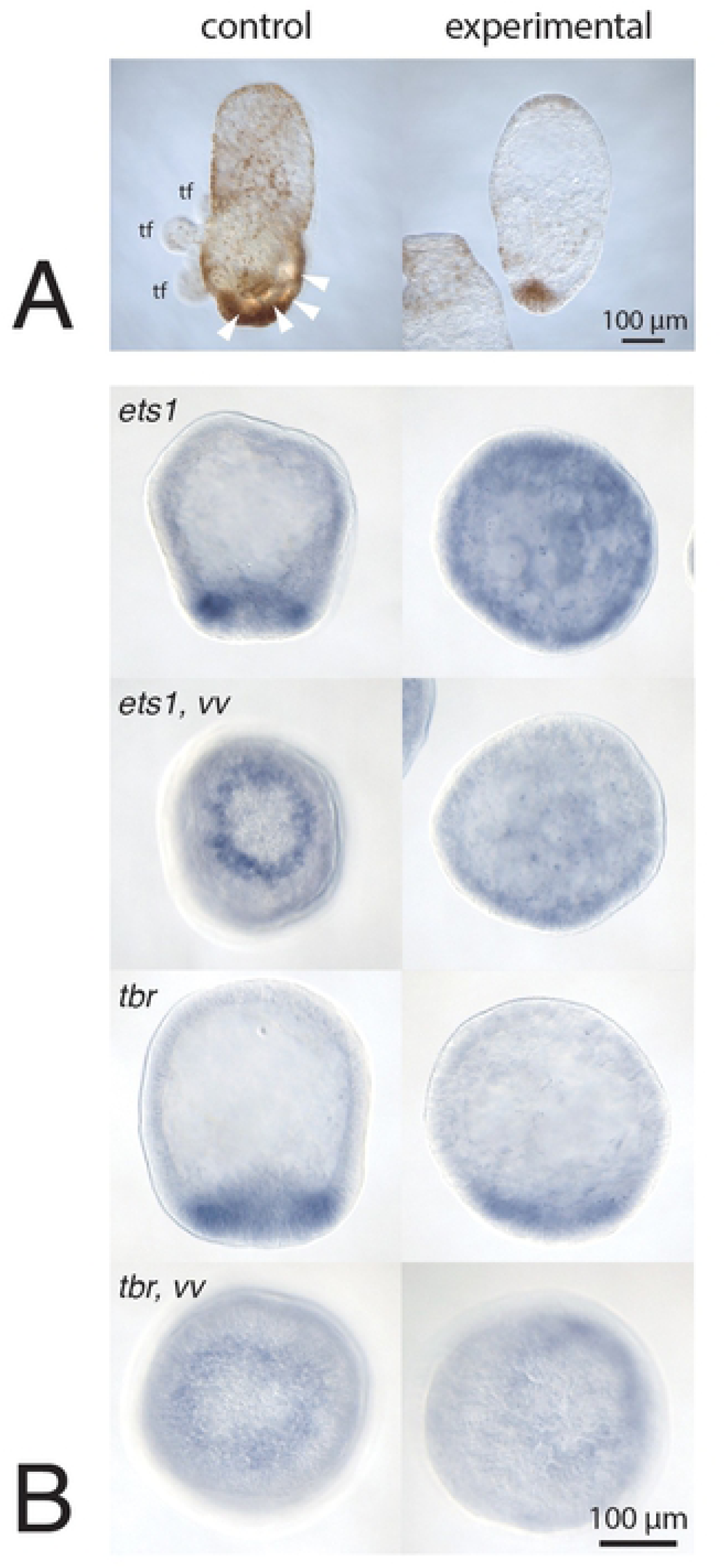
While differentiated cell type phenotypes of the MEK-ERK signal transduction cascade with the small molecule inhibitor UO126 are similar in *H. erythrogramma* as in planktotrophs, the early GRN linkages differ. **A** The MEK-ERK signal inhibitor UO126 treatment reduces skeleton and pigmentation in larva stage *H. erythrogramma* as it does in the ancestral GRN. **B** Despite similar differentiated cell type phenotypes, blastula-stage gene regulatory outputs of MEK-ERK inhibitor treatment in *H. erythrogramma* and the consensus euechinoid. UO126 treatment greatly affects *ets1/2* expression pattern but minimally affects the *tbr* expression pattern.

Thus, regardless of whether the same set of cells coordinate these three signaling pathways in the early embryo, the transcriptional outputs and downstream phenotypic effects of each signaling pathway differ somewhat from the ancestral state, the consensus euechinoid planktotroph GRN.

## Conclusion

### *H. erythrogramma*’s early developmental GRN was rewired to delete the SM-GRN sub-circuit

The sum total of evidence from this study re-casts previous studies in *H. erythrogramma* to suggest a novel conclusion: this species lacks a dedicated larval skeletogenic mesenchyme cell population. In planktotrophic euechinoid sea urchins the SM lineage functions both as the embryonic endomesodermal signaling center and the exclusive source of larval skeletogenic cells in normal development. This cell lineage exhibits 1) unique cell behaviors, such as asymmetric cleavage, early ingression, and directed migration within the blastocoel; 2) a unique suite of co-expressed transcription factors that specify its skeletogenic cell fate, its role as a signaling center, or both; and 3) a defined set of cell signaling interactions by which it induces other endomesodermal cell types and by which its member cells differentiate into skeleton.

Prior studies noted that *H. erythrogramma* lacks a population of cells exhibiting asymmetric cleavage or pre-gastrula ingression [17,87]. Here, we show that it also lacks a population of internalized cells that co-express key larval SM-GRN genes. Taken together, these data suggest that the larval SM lineage as described in indirect developers does not exist in *H. erythrogramma*. Not all echinoderms possess larval skeletons so SM cell identity and signaling center functions clearly do not need to be integrated. In the ancestral euechinoid state, as late-stage larvae approach metamorphosis, skeletogenic cells distinct from larval SM cells and thought to derive from the coelomic pouch mesoderm migrate into the blastocoel and localize near growing larval skeleton [103]. This suggests that prospective juvenile skeletogenic cells are motile, can migrate outside the rudiment, and respond to the same patterning cues as larval SM. We hypothesize that *H. erythrogramma*’s apparent larval skeleton may arise similarly, from cells specified by the juvenile GRN. Interestingly, another echinoderm with independently derived accelerated development retains an unequal cleavage that gives rise to cells that behave similarly to the ancestral SM cells and which do become part (but not all) of the larval skeleton [104]. However, these cells lack the signaling center function [105].

Our results show a surprising degree of re-wiring in the early *H. erythrogramma* gene regulatory network that was not apparent from single-gene or whole-transcriptome studies. Instead, our data suggest the *H. erythrogramma* GRN as a whole is connected differently than previously described GRNs known from sea urchins in which the normal consensus euechinoid SM-GRN is not activated, *i.e.* euechinoid planktotrophs experimentally depleted of SM cells or urchin groups such as cidaroids that specify SM with a different GRN than euechinoids.

We initially considered the hypothesis that *H. erythrogramma*’s mesoderm specification GRN recapitulates a well-documented phenomenon in other sea urchins where experimental removal of precursor or differentiated SM cells triggers activation of the SM-GRN in another cell population to produce replacement SM cells [47,74]. However, *H. erythrogramma*’s GRN does not match this simple model; it is not merely a planktotrophic euechinoid missing SM cells. Co-expression of key skeletogenic markers such as *foxB* and *tbr* is absent from internalized cells, not delayed as in the replacement SM-GRN. In addition, while the MEK-ERK inhibitor UO126 prevents SM (and other mesoderm) specification in *H. erythrogramma* as it does in the ancestral euechinoid, it does not appear to do so by preventing *tbr* transcription, which would be expected for the planktotroph SM-GRN at this stage [106].

We also considered the possibility that the *H. erythrogramma* SM-GRN resembles that of the cidaroid urchin lineage that diverged prior to the evolution of the consensus euechinoid GRN. We found both similarities and striking differences between the two GRNs. Our observation of a Delta signaling input into *alx1* and *hesC* in *H. erythrogramma* resembles the cidaroid GRN [49,51]; however, while cidaroids deploy Tbr in NSM such as pigment cells [51,52] *H. erythrogramma* does not show localized *tbr* expression in any mesenchyme cells. Finally, while some elements appear to be conserved from the ancestral endomesodermal GRN rather than the SM-GRN, such as Wnt and Delta control of *ets1/2* transcription, other connections, such as Wnt signaling activation of *hesC* appear to be *H. erythrogramma* novelties.

**Figure 10.**
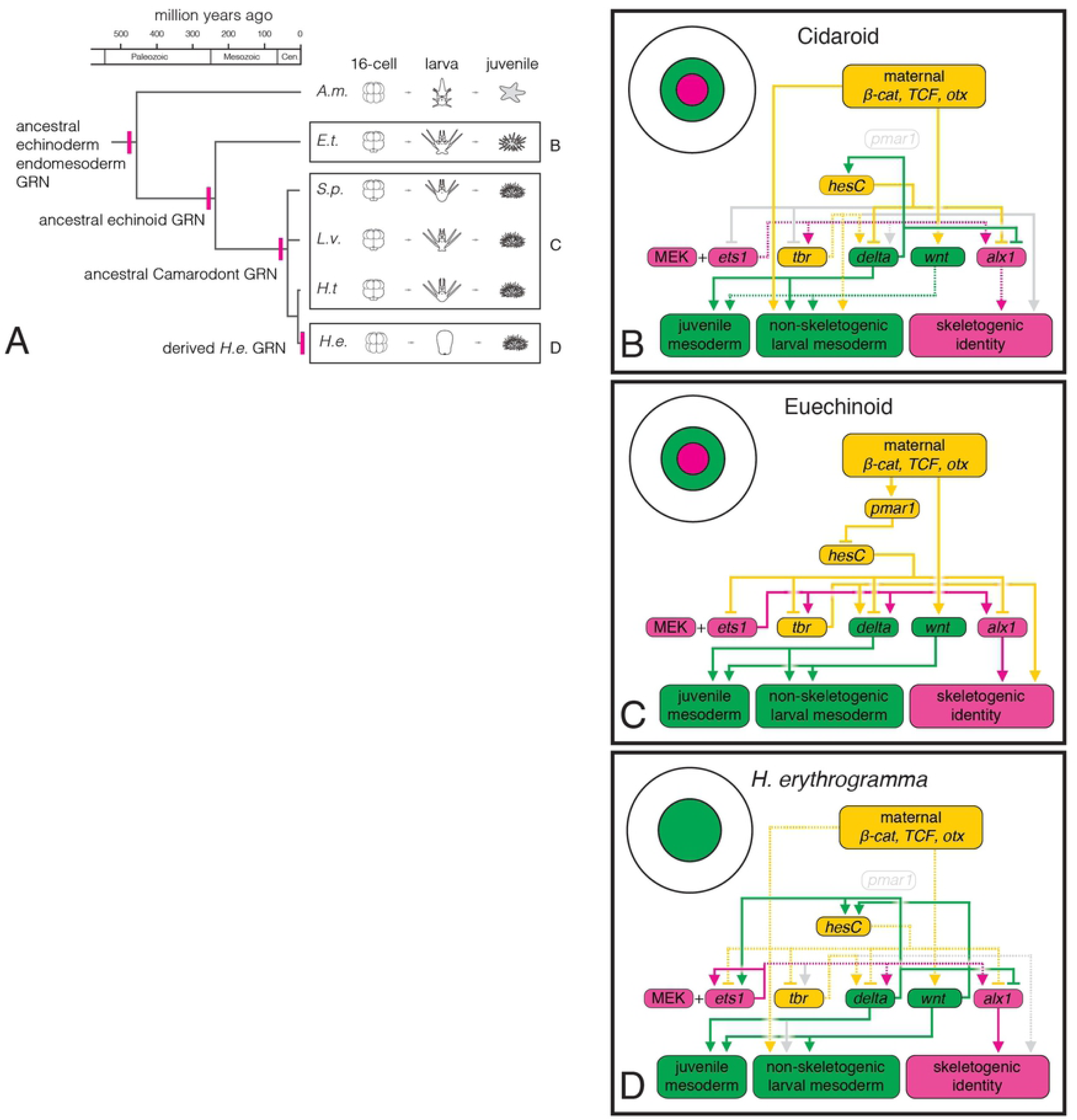
Partial GRN for *H. erythrogramma* mesoderm specification in its evolutionary context. **A** The consensus euechinoid GRN, developed from independent investigations in different sea urchin species, is highly conserved among species diverged ∼40 mya, and many of its features arose with the echinoid GRN 250 mya or earlier. **B** Partial mesoderm GRN for the non-euechinoid echinoids drawn from studies in two cidaroid urchin species, *E. tribuloides* and *P. baculosa*. The consensus euechinoid and non-euechinoid GRNs diverged over >268 mya but show a great deal of conservation, including a MEK-ERK signal requirement for SM and NSM and HesC repression of *alx1*. Features not found in the euechinoid network include repression of *alx1* downstream of Delta signaling and Tbr input into NSM. **C** The consensus euechinoid developmental GRN, also illustrated in Figure 1. Novelties in the euechinoid GRN include the appearance of the double-negative gate logic (Pmar1/HesC double repression) for specifying SM precursors, new HesC regulatory inputs into *ets1/2* and *tbr*, and restriction of *tbr* to the skeletogenic lineage. **D** Our proposed GRN for *H. erythrogramma* mesoderm specification. We found extensive changes to the *H. erythrogramma* GRN despite only ∼4 million years divergence, including loss of the double-negative gate logic for specifying SM precursors, loss of Delta signal induction of non-skeletogenic mesoderm (NSM), and loss of *tbr* from larval SM. In B-D, solid lines show experimentally validated GRN connections. Dashed lines in *H. erythrogramma* and *E. tribuloides* GRNs show connections hypothesized based on indirect evidence such as co-expression or assuming the null hypothesis that they are the same as the euechinoid GRN when no evidence is available. Grey lines in *H. erythrogramma* and *E. tribuloides* GRNs show consensus euechinoid GRN connections absent from those alternative GRNs. Circular diagrams for each GRN represent a vegetal view of fate map for a blastula-stage embryo.

The evolutionary changes in developmental gene expression and cell signaling that we document above are striking in the context of the prior deep conservation of the sea urchin GRN. Many of these features date back at least to the last common echinoid ancestor ∼268 mya and all date back at least to the last common ancestor of the best-studied euechinoids ∼40 mya – yet profound changes have evolved in less than 4 million years within the genus Heliocidaris. Our results indicate that either the long evolutionary conservation of this GRN is not a product of an inherent developmental constraint or that constraint was somehow released. This suggests that even highly conserved features of development, including the earliest steps that pattern the embryo, can be evolutionarily labile under the right conditions. In the case of *H. erythrogramma*, those conditions likely include selection for abbreviated premetamorphic development. We hypothesize that some evolutionary changes to the *H. erythrogramma* GRN, such as removal of the SM sub-circuit described here, are the product of positive selection on interactions within the GRN of early development. Further tests of the GRN to identify stasis or change, formal tests for selection on the genome, and identification of specific cis and trans regulatory changes underpinning GRN differences, and similar studies in other lecithotrophic urchins will help to identify points of lability and constraint in the developmental GRN.

## Methods

### Reagents

Reagent brand and stock information is detailed in Supplemental File 1.

### Animals and embryo cultures

Adult *H. erythrogramma* and *H. tuberculata* were obtained off the east coast of Australia at Little Bay, New South Wales (33°58’S, 151°14’E) and maintained in natural sea water aquaria at ambient temperature (20–23°C). Adult *L. variegatus* were collected near Duke University Marine Lab in Beaufort, NC USA (34°43’N, 76°40’W) or obtained commercially from Reeftopia (Key West, FL, USA) and maintained in artificial seawater at ambient temperature (20–23°C). Animals were spawned by intracoelomic injection of 0.5 M KCl and gametes collected in Millipore-filtered natural sea water (FSW). Control time course embryos were cultured in FSW. Embryo cultures were maintained at ambient temperatures or in a cooling water bath set at 22°C. Time points are summarized in Table 1, detailed version in Supplemental File 1).

**Table 1:** Key stages in H. erythrogramma development at ∼22° C

| stage | hpf | key features |
| --- | --- | --- |
| unfertilized egg | 0 | egg |
| wrinkled blastula | 6-8 | early blastula |
| **hatched blastula** | 10-12 | embryo hatches from fertilization envelope |
| early gastrula | 18 | archenteron begins invagination |
| mid-gastrula | 24-26 | archenteron full-length |
| early larva | 32 | coelom compartments; vestibule ingression |
| early rudiment larva | 36 | skeleton biomineralization begins |
| **late rudiment larva** | 52-56 | larval and juvenile skeletal elements co-occur |
| early metamorphosis | 72-120 | tube feet emerge; extensive juvenile skeleton |

### Morpholine-substituted oligonucleotides

MASOs were designed against the translation start sites of target genes and synthesized by Gene Tools. MASO sequences and effective concentrations are in Table 2. Morpholino doses were titrated empirically to the lowest effective dose.

**Table 2:**
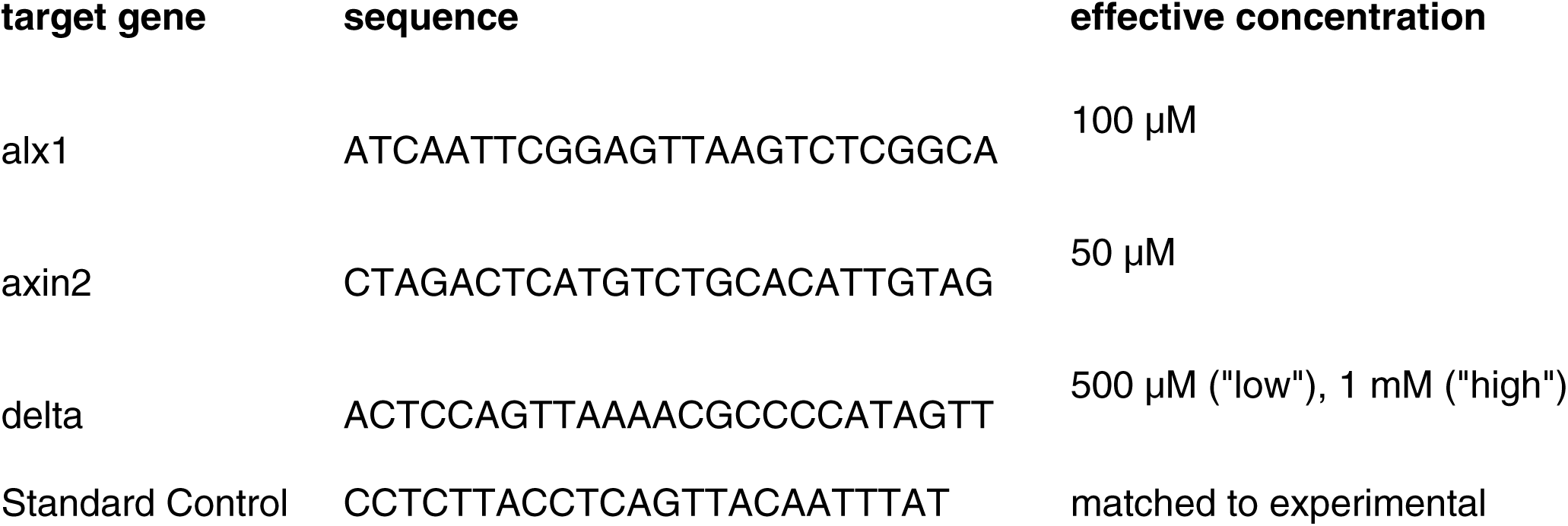
Translation-blocking MASO sequences

### Microinjection

Microinjection was performed as described in (Edgar et al, in review). Needles were pulled on a Sutter p97 micropipette puller from WPI needle stock (TW100F-6). Reagents were mixed with fluorescent injection mix (RNase-free 2X injection mix: 3.5 μl water, 6.5 μl 150 mg/ml lysine-fixable fixable TMR dextran 10,000 MW, 2.0 μl 4M KCl, 8.0 μl glycerol). Fertilized embryos were injected before first cleavage on agarose pads in a solution of pasteurized (30 minutes 65°C) filtered seawater (PFSW) + 2% w/v Ficoll 400 (Sigma F-9378). Embryos were hand-sorted for fluorescence between second and sixth cleavage cycles. Injected embryos were cultured in IVF dishes (Thermo-Fischer 176740) or gelatin-coated dishes in PFSW + penicillin (100 unit/ml) and streptomycin sulfate (0.1 mg/ml) (Sigma P4333A).

### Fixation

For general morphological analysis, ISH, and IHC, embryos were fixed overnight (∼16 hours) at 4°C in 4% paraformaldehyde (Sigma 158127) + 20 mM EPPS (Sigma E1894), washed 3 times in pasteurized filtered seawater, and dehydrated step-wise into 100% methanol and stored at −20°C in non-stick tubes.

For biomineralized skeleton morphological analyses, embryos were fixed with 2.5% (v/v) glutaraldehyde (ProSciTech, Australia) in filtered seawater for 1 hour at 4°C, washed in FSW, dehydrated in an ethanol series to 70% (v/v) ethanol in Milli-Q water, adjusted to pH 7.8 with glycerophosphate (after [112,113] and stored at −20°C. To image, embryos were dehydrated completely into methanol and cleared in 2:1 (v/v) benzyl benzoate: benzyl alcohol.

### Probe constructs

PCR primers were designed from mRNA sequences in the reference transcriptome published in [14] using PrimerBLAST (NCBI) and synthesized by IDT or EtonBio. Primer sequences are listed in Table 3.

**Table 3:**
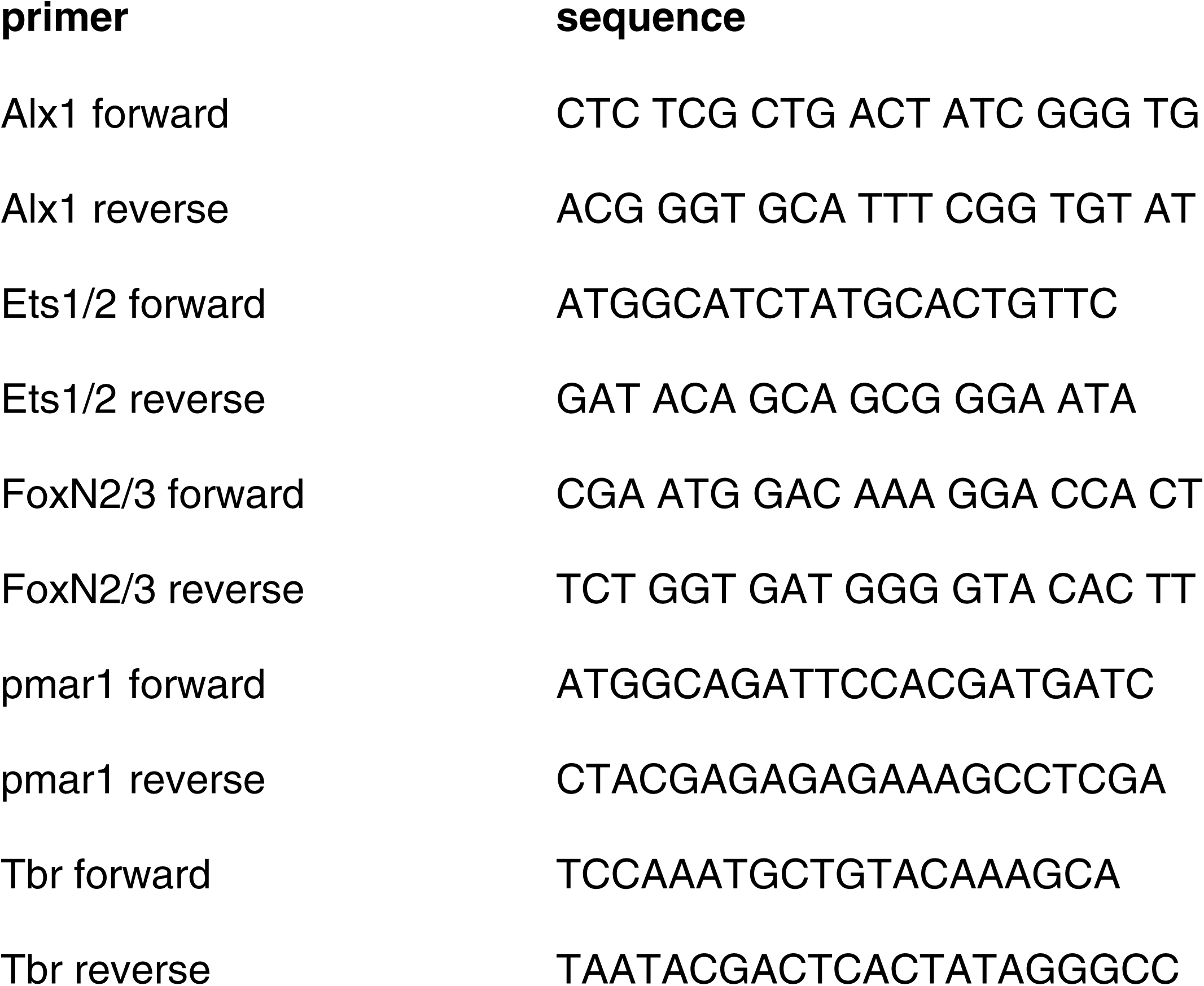
Primers

Probe inserts were ligated into pGEM T-easy (Promega) according to kit instruction. Full-length He-HesC was synthesized in vitro by GenScript and subcloned. Plasmid information is in Table 4.

**Table 4:** Plasmids

| **name** | **insert NCBI number** | **reference** |
| --- | --- | --- |
| He-Alx1-in-pGEM-T | MK749160 | this study |
| He-Ets1-probe-pGEM-T | MK749161 | this study |
| He-Tbr-probe-pGEM-T | MK749162 | this study |
| He-HesC-FL-in-pBS | MK749159 | this study |
| He-Delta |  | Koop et al 2017 |
| He-FoxN2/3-probe-pGEM-T | MK749163 | this study |
| He-FoxB-probe-pGEM-T |  | this study |
| He-pmar-probe-pGEM-T | MK876229 | this study |

### Small molecule inhibitor treatments

Small molecule effective doses were empirically titrated with starting doses above and below published effective concentrations for other echinoderms; optimal doses were close to published values from other sea urchin species. Effective concentrations are in Table 5. For scored treatments and ISH analysis, biological replicates consisted of 3 unique crosses, typically fertilized, treated, and fixed in parallel to ensure similar ambient temperatures (however, DAPT time course experiment, Supplemental Figure 1A, includes fewer biological replicates; raw data in Supplemental File 1). Vehicle controls were treated with an identical volume of the same solvent. We chose to score for presence/absence of skeletal elements because the size and number of elements may be affected independently of initial specification, while inhibitors tested may have effects on adult skeletogenic cells; for example, DAPT is known to inhibit differentiation of adult sea urchin skeletogenic cells [107].

**Table 5:** Small molecule inhibitors

| inhibitor | concentrations tested | optimal dose | vehicle |
| --- | --- | --- | --- |
| DAPT | 5, 8, 10, 12, 16 μM | 5 μM ("low"), 10 μM ("high") | DMSO |
| LY-411575 | 0.1, 1.0, 10 μM | 1 μM | DMSO |
| UO126-EtOH | 6, 12, 24 μM | 12 μM | DMSO |
| C59 | 0.5, 1.0, 2.0 μM | 2.0 μM | DMSO |
| LiCl | 20 mM | 20 mM | H<sub>2</sub>O |

### Whole-mount in situ hybridization

Chromogenic whole mount in situ hybridization after the methods previously published [23] and detailed in Table 6. Briefly, digoxygenin-labeled RNA probes were prepared from either restriction-digested plasmids or PCR products containing a T7 promoter site. Control ISH patterns were determined using a mix of at least 3 biological replicates (control cultures from unique crosses). Hybridizations were carried out at 65°C and stringency washed at 0.1% SSC.

*H. erythrogramma* to *H. tuberculata* comparison ISH were carried out in parallel with *H. erythrogramma* probes (*ets1/2, foxB, foxN2/3, hesC, pmar1, tbr*) using the same reagents and equipment; each included 2-3 biological replicates for each developmental stage. Sense probes prepared from the same constructs and no probe controls did not exhibit localized expression patterns.

### Whole-mount immunohistochemistry

Immunohistochemistry protocol is summarized in Supplemental File 7. The Endo-1 monoclonal antibody labels sea urchin endoderm [114] (used at 1:100, mouse IgG) and 1D5 recognizes the skeletogenic cell-specific cell-surface protein msp130 [115] (used at 1:50, mouse IgM). Secondary antibodies (goat-anti mouse IgG and IgM conjugated with AlexaFluor 647, 488) were used at 1:1000. Hoescht was used at 1:10,000 to counterstain nuclei.

### Image capture

*H. erythrogramma* embryos were washed with 100% ethanol or methanol, then were cleared and mounted in 2:1 (v/v) benzyl benzoate: benzyl alcohol (BB:BA). *H. tuberculata* embryos were cleared with either BB:BA or 50% glycerol. *L. variegatus* embryos were cleared with 50% glycerol. DIC and fluorescence micrographs were taken on either an Olympus BX60 upright microscope with an Olympus DP73 camera or a Zeiss Upright AxioImager with a Zeiss MRm or a Zeiss ICc1 camera using ZEN Pro 2012 software.

### Image manipulation and scoring

Morphological measurements were made in ImageJ 2.0.0 using the standard Measure tool. Presence/absence measurements were scored manually. Fixed samples were viewed under polarized light to visualize the birefringent calcite skeleton, and under white light to visualize pigment cells and general morphology. Larval and prospective juvenile skeletal elements are identified by morphology: larval elements are bilaterally symmetrical according to the larval ectoderm while juvenile elements are arranged in a pentamerally symmetric pattern in a plane on the prospective oral juvenile ectoderm.

Illustrations were drawn, figure panels were assembled and additions such as arrows, panel labels, and scale bars were added with Adobe Illustrator. No other adjustments were made except to the fluorescent images (Figure 3D), which were contrast-adjusted using identical cutoff values in ZEN Pro to reduce background fluorescence. Raw .czi files are available as Supplemental Files 2 and 3.

### Gene expression analysis

We analyzed gene expression of key skeletogenic and endomesoderm GRN genes based on a previously published data set [14] using the R packages edgeR 3.16.5 [116] and maSigPro 1.46.0 [117]. An R Markdown file to replicate these results is available in Supplemental File 4 (requires Table S6 of Israel et al 2016, journal.pbio.1002391.s015.csv, as input).

## Acknowledgements

Several former members of the Byrne lab contributed materially to this project. Thanks to Demian Koop for acquiring the images in Figure 2 A-F and rearing and collecting some of the *H. tuberculata* embryos, Paula Cisternas for collecting preliminary data (not shown) with the MEK inhibitor UO126, and both of them for providing lots of advice on *H. erythrogramma* protocols. Lingyu Wang treated and fixed all the embryos in Figure 2 as well as some control time course embryos. We gratefully acknowledge Matt Naylor, Pauline Aubel, and Haydn Allbutt who kindly provided access to key pieces of equipment at the University of Sydney. Steven D. Black provided insightful feedback on a previous version of this manuscript, as did many members of the Wray and McClay labs.

This work was funded by National Science Foundation grant IOS-1457305, by Australian Research Council grant DP120102849, and by NIH RO1 HD14483 and PO1 HD37105.

## Supplemental Figures

**Supplemental Figure 1.**
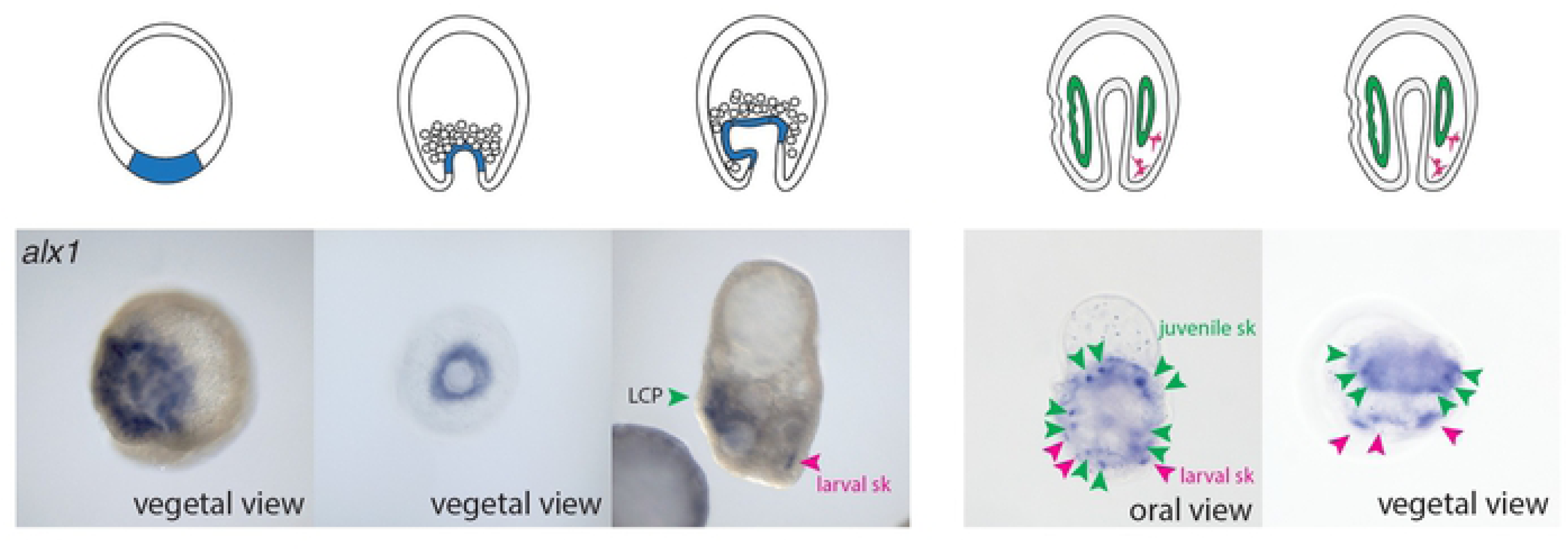
Spatiotemporal expression of *alx1* in *H. erythrogramma* (supplemental to Figure 2). External, vegetal view at hatched blastula stage shows *alx1* expression throughout the vegetal pole rather than in a ring. At early gastrula stage, vegetal view shows *alx1* restricted to the archenteron. From late gastrula, *alx1* is expressed in the left coelomic pouch (green arrowheads) but not overlying ectoderm. Additional *alx1* expression at the site where vestigial larval skeleton will be synthesized (pink arrowheads) is the earliest demonstrated localized expression of any SM marker at the prospective site of larval skeletogenesis in *H. erythrogramma*. Foci of juvenile *alx1* expression arranged in a pentamerally symmetrical pattern with two foci per tube foot (green arrowheads) as well as larval *alx1* expression behind the rudiment (pink arrowheads).

**Supplemental Figure 2.**
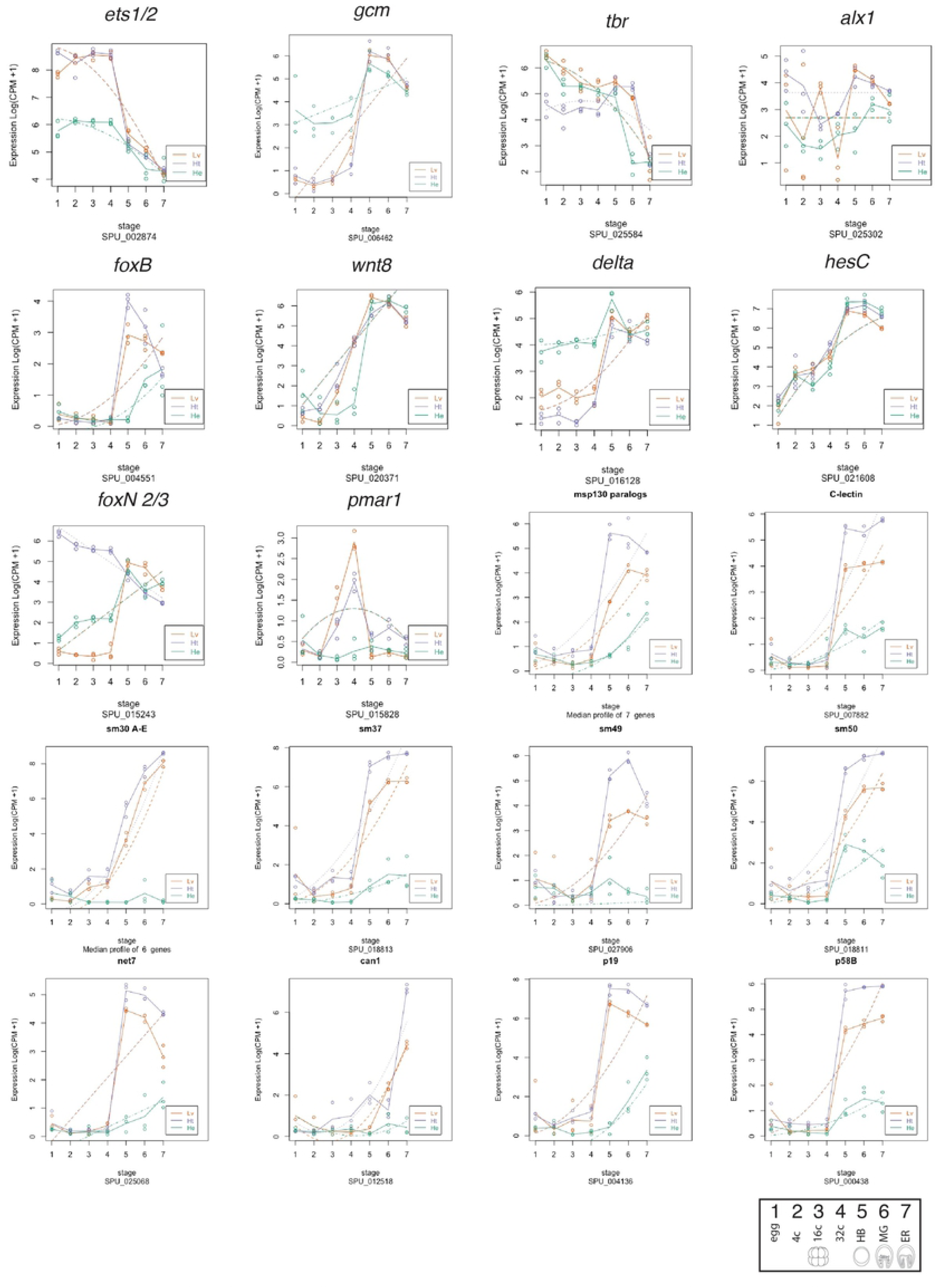
Expression profiles of key SM genes in from whole-transcriptome profiling in *H. erythrogramma, H. tuberculata* and *L. variegatus* at equivalent stages. Interestingly, in contrast to *ets1/2*, which is expressed at a much lower level in early *H. erythrogramma* embryos, *delta* and its target gene *gcm* are expressed at a higher level. These early NSM genes each show quantitative differences in expression in *H. erythrogramma* although the timing of their expression changes resembles planktotrophs. Note that skeletogenic differentiation genes (K-T) are all significantly delayed in *H. erythrogramma*.

**Supplemental Figure 3.**
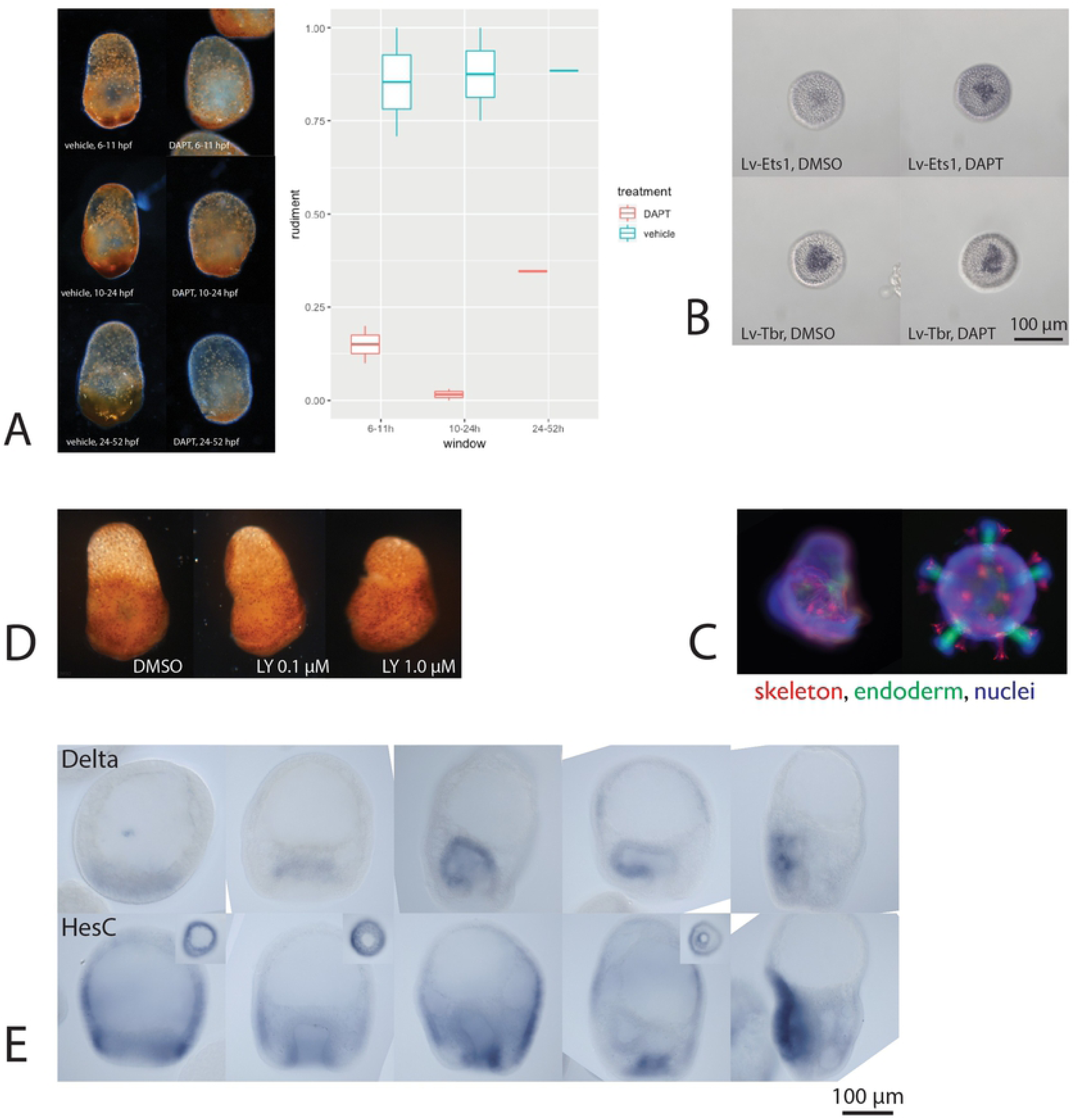
**A** Inhibition of Delta/Notch signaling by DAPT does not eliminate pigment cells at any time point but affects coelomic pouch specification throughout early development. Coelomic pouch specification is strongly inhibited any time prior to or during gastrulation. Raw data in Supplemental File 1. Note that reduced biomineralization of skeleton with DAPT treatment has been observed previously in other sea urchins [107], so raw counts of biomineralized elements do not reflect presence/absence of skeletogenic cells. **B** The gamma-secretase inhibitor LY411575 is an even more specific inhibitor of Delta/Notch signaling than DAPT [108,109], which has some off-target effects in the p38 MAPK pathway[110,111]. Even high doses of the inhibitor do not eliminate pigment cells. **C** DAPT inhibitor treatments in the indirect developing urchin *L. variegatus* confirm results as predicted by knockdown experiments in several species used to construct the consensus indirect developer GRN (DAPT’s effect on *hesC* and *delta* expression in a model indirect-developing euechinoid [92]). **D** Delta and HesC mRNAs are expressed in complementary patterns during much of *H. erythrogramma* development.

